# Unusual intragenic suppression of an IFT52 gene disruption links hypoxia to the intraflagellar transport in *Tetrahymena thermophila*

**DOI:** 10.1101/044420

**Authors:** Drashti Dave, Gautham Pandiyan, Dorota Wloga, Jacek Gaertig

## Abstract

IFT52 protein is a conserved intraflagellar transport protein (a part of the IFT complex B) that is essential for assembly and maintenance of cilia. *Tetrahymena* null mutants with an insertion of a *neo* gene cassette into the *IFT52* gene undergo frequent suppressions that lead to conditional assembly of cilia only under hypoxic conditions (Brown et al. 2003). Here we show that these conditional suppressions are intragenic and occur by a novel mechanism. First, the non-native (bacterial) portion of the DNA sequence of the *neo* cassette is deleted during the process of genome rearrangement that occurs in the developing macronucleus of conjugating *Tetrahymena*. Next, the residual sequences of the *neo* cassette (of *Tetrahymena* origin) within the IFT52 mRNA are recognized as multiple introns and undergo splicing, leading to a restoration of the translational frame of *IFT52*. The resulting hypoxia-dependent IFT52 protein contains an insertion of 43 new amino acids that replace 7 original amino acids. Taken together with a study in *Chlamydomonas reinhardtii* showing a hypoxia-dependence of another IFT subunit mutant, IFT46, (Hou et al. 2007), our observations generalize that defective IFT complex subunits can regain functionality under hypoxia.

## Results and Discussion

Intraflagellar transport (IFT) is a bidirectional motility of ciliary precursors that occurs inside cilia (Kozminski et al. 1993). Kinesin-2 is the anterograde IFT motor, whereas cytoplasmic dynein1b is responsible for the retrograde IFT (Kozminski et al. 1995; Pazour et al. 1999; Porter et al. 1999). These motors move IFT trains, that are composed of two protein complexes, A and B (Cole et al. 1998; Piperno and Mead 1997). IFT52 is a complex B protein that is required for the assembly and maintenance of cilia (Brazelton et al. 2001; Brown et al. 2003; Deane et al. 2001).

The ciliate *Tetrahymena thermophila* has two nuclei, a transcriptionally silent micronucleus (Mic) and a transcriptionally active macronucleus (Mac) (reviewed in (Yao and Chao 2005)). Earlier (Brown et al. 2003), *IFT52* was disrupted by insertion of the *neo2* marker within exon 4 (Figure 1D). Heterokaryons were constructed with Macs carrying wild-type *IFT52* alleles and Mics homozygous for the disrupted alleles. Most progeny cells of mating *IFT52* heterokaryons are completely paralyzed due to the lack of cilia and cannot complete cytokinesis since they are unable to rupture the connecting cytoplasmic bridge (rotokinesis). Surprisingly, 3% of the heterokaryon progeny recover partial motility due to spontaneous suppressions. Importantly, the suppressed cells (IFT52Δsm) assemble motile cilia when grown at either a lower temperature or in hypoxia. In a single suppressed strain, an additional event produced IFT52Δmov cells, which are capable of assembling cilia independently of temperature or hypoxia (Brown et al. 2003).

**Figure 1.**
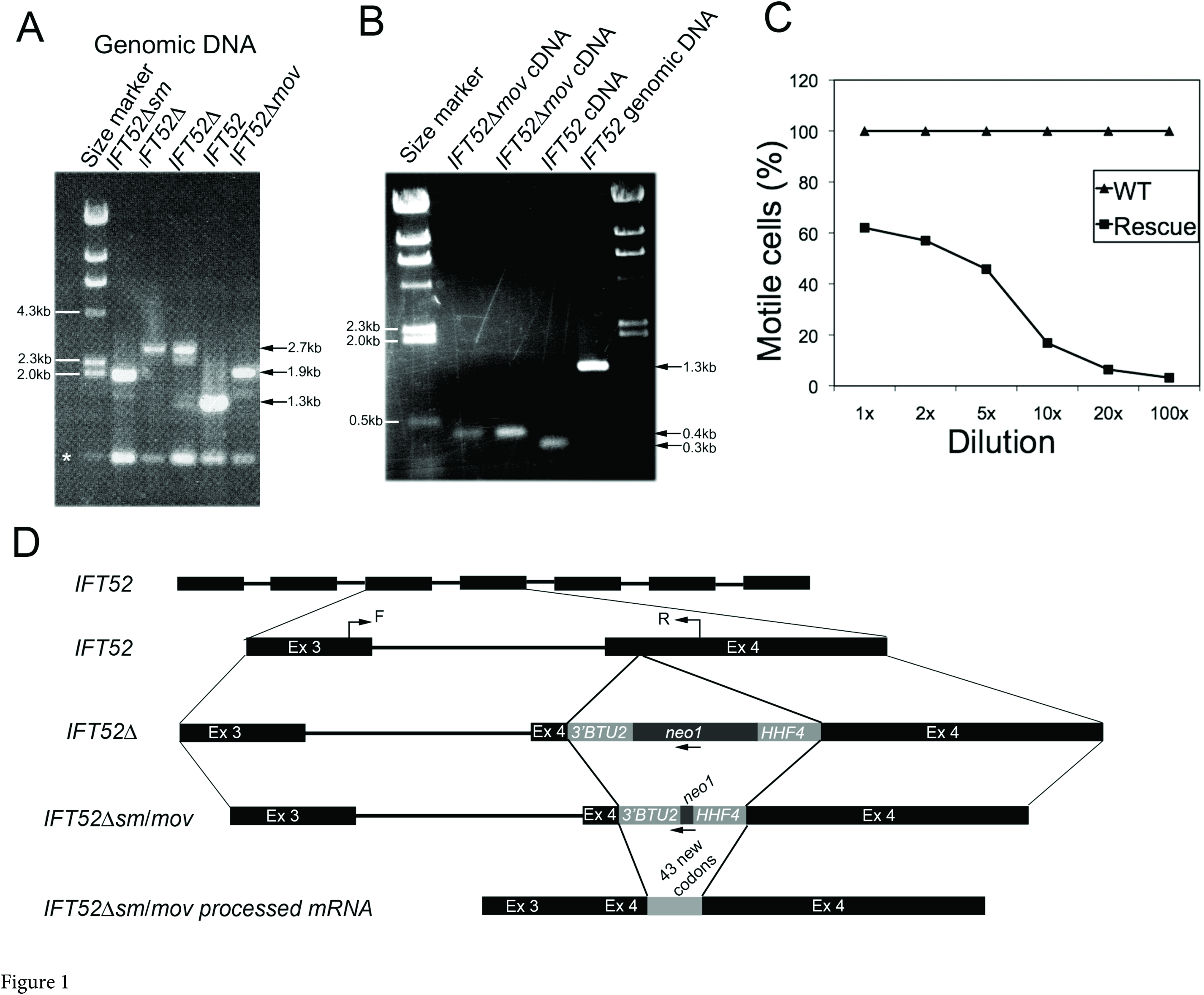
Two subsequent sequence deletions lead to intragenic suppression of an insertional *IFT52* mutation. A. Results of PCR amplifications of the genomic region of IFT52 locus with primers corresponding to sequences in exons 3 and 4 in wildtype (*IFT52*), gene knockout (*IFT52Δ*) and the suppressor (*IFT52Δsm/mov*) cells. Amplified fragments were separated on an agarose gel. An asterisk marks an apparent non-specific amplification product. B. PCR amplifications of total cDNA obtained from mRNA using wildtype (*IFT52*), knockout (*IFT52Δ*) and suppressors (*IFT52Δsm/mov*) cells using the same primers as in panel A. The amplification products were separated on an agarose gel. C. IFT52Δ cells rescued with *IFT52Δsm/mov* DNA and wild-type cells both at a concentration of 3x10^^5^ were diluted down to different concentrations (1x to 100x) on a 96-well microtiter plate, and incubated at 30°C. After 12 hours of incubation, the number of motile cells (cells showing detectable displacement and lacking cytokinesis defects) was determined. D. A schematic diagram detailing the *IFT52* locus in the wild-type (*IFT52*), knockout (*IFT52Δ*) and the suppressors (*IFT52Δsm/mov*) cells and the cDNA in the suppressors (*IFT52Δsm/mov processed mRNA)* cells. F and R represent primers used for the various PCR reactions.

The high frequency of the IFT52Δsm conditional suppressions and the fact that these suppressions occur only during conjugation (Brown et al. 2003), suggested that the mechanism of suppression is based on processes that occur inside the developing new Mac. Conjugating *Tetrahymena* cells undergo a series of nuclear events that culminate in replacement of the parental Mac by a new Mac that develops by differentiation from a zygotic Mic (reviewed in (Coyne et al. 1996)). About 15% of the Mic genome is removed from the new Mac, by a pathway that involves an RNAi-dependent sequence recognition and degradation (reviewed in (Yao and Chao 2005)). Yao and colleagues showed that a foreign sequence, *neo2*, inserted into multiple loci, undergoes RNAi-mediated deletion (Yao et al. 2003). Thus, we tested whether *neo2* inserted into *IFT52* also undergoes deletions that could be a cause of the conditional suppressions.

The *IFT52* knockout was done by inserting the *neo2* disruption cassette into exon 4 (Figure 1D). *neo2* consists of the bacterial neomycin phosphotransferase (*neo*) coding region placed between DNA fragments of *Tetrahymena* origin; the *HHF4* promoter and the *BTU2* transcription terminator (Gaertig et al. 1994; Kahn et al. 1993). We isolated total genomic DNA from wild-type, IFT52Δ, IFT52Δsm and IFT52Δmov cells and amplified the *IFT52* locus across the *neo2* insertion site (Figure 1D). Amplification of genomic DNA of wild-type cells produced a fragment of expected size (1.3kb). The same primers used with IFT52Δ (non-suppressed) DNA produced a larger fragment (~2.7kb) consistent with presence of an intact *neo2* cassette (Figure 1A). Strikingly, the same primers amplified a smaller fragment (~1.9kb) from the genomic DNA of both conditional and non-conditional suppressors (IFT52Δsm and IFT52Δmov). This suggested that the suppressions are associated with deletions around the *neo2* insertion site. Sequencing of fragments amplified from multiple independent suppressor strains showed deletions of a portion of *neo* (~0.8kb) with deletion junctions at exactly the same positions, while the flanking sequences of *neo2* (of *Tetrahymena* origin) remained largely intact (Figure 1D). Specifically, all deletions analyzed were between the nucleotide at position +45 in the *neo* coding sequence and the fifth nucleotide downstream of the stop codon within the *BTU2* segment (Figure S1). These observations are consistent with earlier reports on deletions of *neo2* sequences during macronuclear development (Liu et al. 2005; Yao et al. 2003).

The observed *neo* sequence deletions, do not explain the mechanism of suppression because the sequence of the *neo2* cassette remnant has stop codons in all forward translational frames. An Ift52p translated from the predicted mRNA containing the *neo2* remnant would be severely truncated; lacking 5 out of 7 exons, all containing conserved sequences (Cole 2003) (Figure S1). Nevertheless, the suppressions correlate with deletions of the *neo* coding sequence.

To establish whether the suppressed *IFT52* locus with the residual *neo2* is sufficient to restore partial motility, we introduced the rearranged fragment of the *IFT52Δmov* genomic DNA into IFT52Δ cells by biolistic bombardment (Figure 1D). As a control we mock-transformed the same number of IFT52Δ cells (9x10^6^). After 7-9 days of incubation at room temperature, we obtained 2 clones that regained motility in the population bombarded with the rearranged (*IFT52Δmov)* fragment and none in the mock-transformed IFT52Δ cells. We confirmed that the targeting fragment replaced the corresponding region of the fully disrupted *IFT52* locus by PCR (results not shown). The rescued cells showed the conditional suppression phenotype, a cell density (pericellular hypoxia)-dependent ciliary motility (Brown et al. 2003) (Figure 1C and results not shown). These data indicate that the IFT52Δmov cells underwent an additional, unknown genetic or epigenetic change that resulted in a non-conditional suppression.

The rearranged *IFT52Δsm/mov* gene contains a residual *neo2* sequence that somehow provides a partially functional Ift52p. Either an extremely truncated Ift52p is sufficient for conditional ciliary assembly or an additional mechanism restores the translational frame across the residual *neo2*. To determine the sequence of the translated Ift52p in IFT52Δsm cells, we used RT-PCR to amplify the *IFT52* cDNA obtained from mRNA of IFT52Δsm cells (Figure 1D). For a spliced wild-type *IFT52* mRNA, the amplified fragment was expected to be ~0.3kb. For the *IFT52Δsm* mRNA with residual *neo2* cassette, the cDNA fragment was expected to be ~0.9kb. However, the size of the amplified product from the IFT52Δsm cDNA was ~0.4kb, indicating that an additional splicing event occurs in the IFT52Δsm mRNA (Figure 1B). The sequencing of a cloned IFT52Δsm cDNA revealed that ~0.8kb of the residual *neo2* was absent. Most of the residual *neo2* sequence, mainly comprising of the *HHF4* and *BTU2* sequences, was removed from the mRNA as 3 (artificial) introns. The artificial intron junctions have sequences consistent with the native intron junctions observed in ciliates such as *Paramecium* and *Tetrahymena* (Figure S1) (Jaillon et al. 2008). The processing of the residual *neo2* as a set of artificial introns restores the translational frame across the site of *neo2* insertion (Figure S2). Hence, the predicted suppressor Ift52p has 43 additional amino acids but lacks 7 original amino acids as a result of the *neo2* cloning procedure (Figures 1D and S2). Either the presence of these extra amino acids or the absence of the 7 endogenous amino acids in Ift52p-sm (or both) results in the intragenic conditional suppression.

To conclude, we reveal a novel mechanism for intragenic suppression in *Tetrahymena* that consists of two steps: 1) foreign DNA within the inserted disruption cassette is deleted during macronuclear development, and 2) the remaining AT-rich *Tetrahymena* native sequences of the disruption marker are processed as introns during mRNA splicing. The first step almost certainly occurs via the RNAi-mediated developmental genome rearrangement pathway (Mochizuki et al. 2002; Mochizuki and Gorovsky 2004; Yao et al. 2003). This form of genomic DNA deletion is thought to have evolved as a means of genome surveillance to eliminate transposon DNA from the transcriptionally active Mac (reviewed in (Yao and Chao 2005)). In 4 independent suppressor clones we detected a genomic deletion at precisely identical positions. Previous studies showed a variability in the deletion sites (Liu et al. 2005; Yao et al. 2003). It is likely that other deletions occur in the disrupted *IFT52* locus but they do not create potential splice junctions that restore the translational frame. When *IFT52* heterokaryons undergo conjugation, the majority (97%) of the progeny has a non-suppressed phenotype. In these cells, the deletions of *neo2* either do not occur, or occur on an insufficient number of macronuclear chromosomes to achieve a phenotypic threshold for suppression (there are 45 copies of each chromosome in the G1 macronucleus).

*Chlamydomonas* cells carrying an insertional mutation in IFT46 (encoding another complex B protein), also underwent a spontaneous intragenic mutation that led to a hypoxia-dependent cilia assembly (Hou et al. 2007). Both studies taken together ((Hou et al. 2007) and this work), allow for a generalization; that hypoxic conditions can restore the functionality of mutated IFT complex B components. Hou and colleagues observed the assembly of complex B in the flagella of suppressed *Chlamydomonas* IFT46 mutants. It is likely that the suppressed IFT52Δsm *Tetrahymena* cells also assemble complex B. Hou and colleagues proposed that the IFT complex B subunits are folded by a chaperone whose levels increase under hypoxia. Thus, a partly damaged IFT component may still fold properly when the chaperone activity is increased. Another possibility is that the IFT complex B assembly is regulated directly by an oxygen-dependent post-translational modification of one or more subunits. Regardless of the exact mechanism, both studies indicate that a hypoxia-dependent modulation of the activity of IFT complex B subunits is a conserved mechanism.

## Materials and Methods

### Cells, cultures and media

For the maintenance, IFT52Δ, IFT52Δsm and IFT52Δmov cells were grown at the room temperature in MEPP medium (Orias and Rasmussen 1976) with an antibiotic-antimycotic mixture (Invitrogen, Carlsbad, CA).

### DNA preparation and cloning

Isolation of total genomic DNA was done as described (Dave et al. 2009). The genomic region across exons 3 and 4 was amplified using the following primers: 5’-ATGCCCTCAAATAAT-3’ and 5’-TAGAGTTGGTTTAGATTT-3’. The resulting fragments were cloned into pGEM-T-vector (Promega Corp, Madison, WI) and sequenced.

### Biolistic transformation of Tetrahymena

To determine whether a genomic fragment of IFT52 of suppressor origin is sufficient to confer suppression of the IFT52Δ phenotype (lack of cilia), IFT52Δ cells were biolistically bombarded with a genomic fragment of IFT52Δmov origin, that was earlier separated from the pGEM-T-vector plasmid with NcoI and SalI digestion. Bombarded cells were grown at the room temperature and transformants were identified based on recovery of cell motility (Cassidy-Hanley et al. 1997).

### cDNA preparation

Cells were grown to a concentration of 2 x 10^5^ cells/ml in MEPP medium (Gorovsky et al. 1975), washed with 10 mM Tris-HCl buffer pH 7.5 and used for total RNA extraction with TRI-reagent (MRC Inc, Cincinnati, OH) according to manufacturer’s instructions. Total cDNA was prepared using the SMART IV-forward and CDS III-reverse primers from the RT-PCR kit (Clontech Inc, Mountainview, CA).

## Acknowledgements

This work was supported by the National Science Foundation grant MCB-063994 to JG.

## Supplemental data

**Figure S1.**
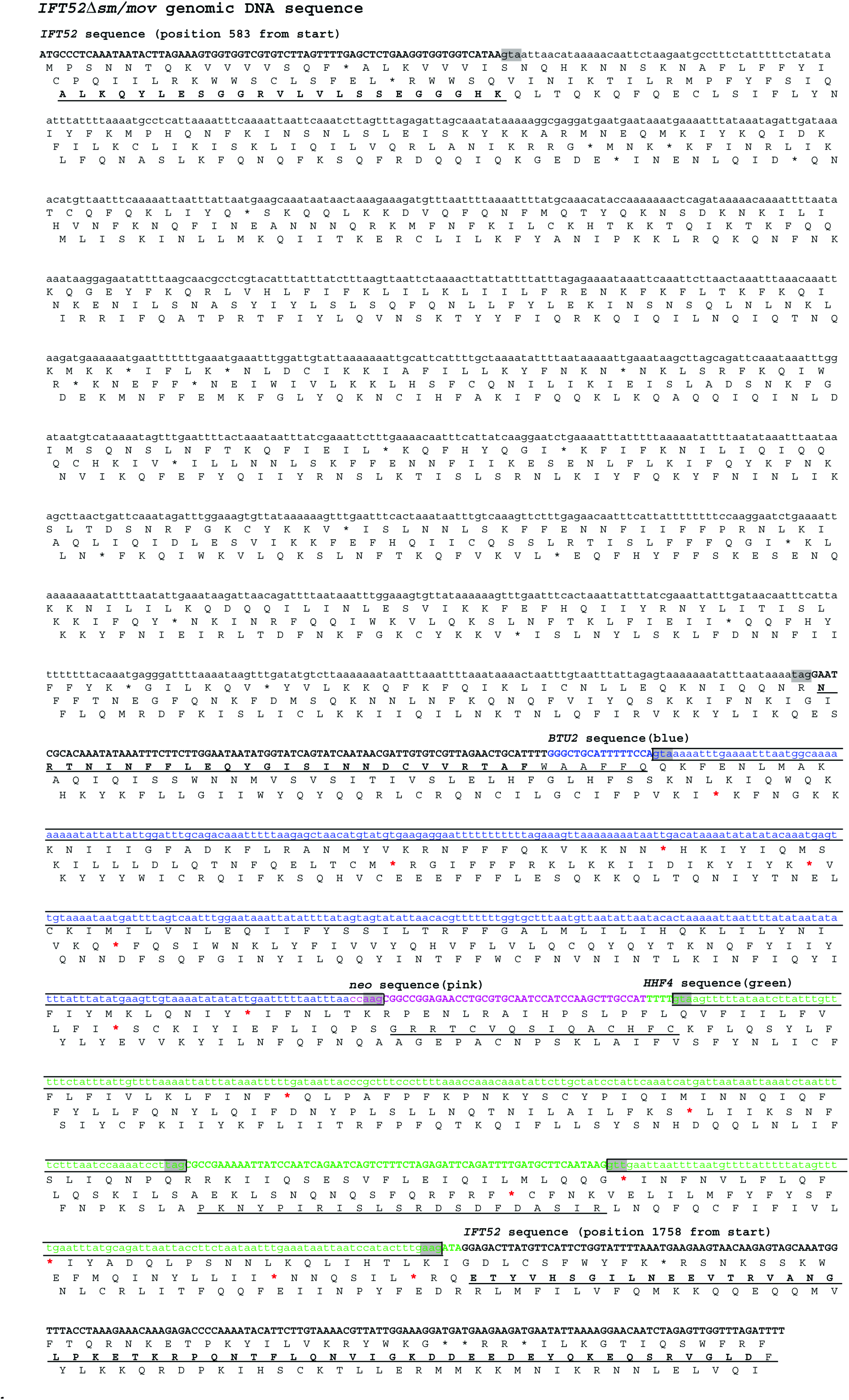
Sequence of genomic DNA reconstructed from 4 independent IFT52Δsm/mov suppressor strains shows a deletion within the *neo2* cassette. The sequence shown corresponds to the region between exons 3 and 4 of the suppressed *IFT52* genomic DNA. The three segments of the *neo2* cassette are marked in blue (*BTU2*), pink (*neo*) and green (*HHF4*). Natural intron junctions are marked as grey boxes. Artificial intron junctions are marked as open boxes. Note: Within the residual *neo* cassette left after genomic deletion, there are stop codons (red *) in every translational frame.

**Figure S2.**
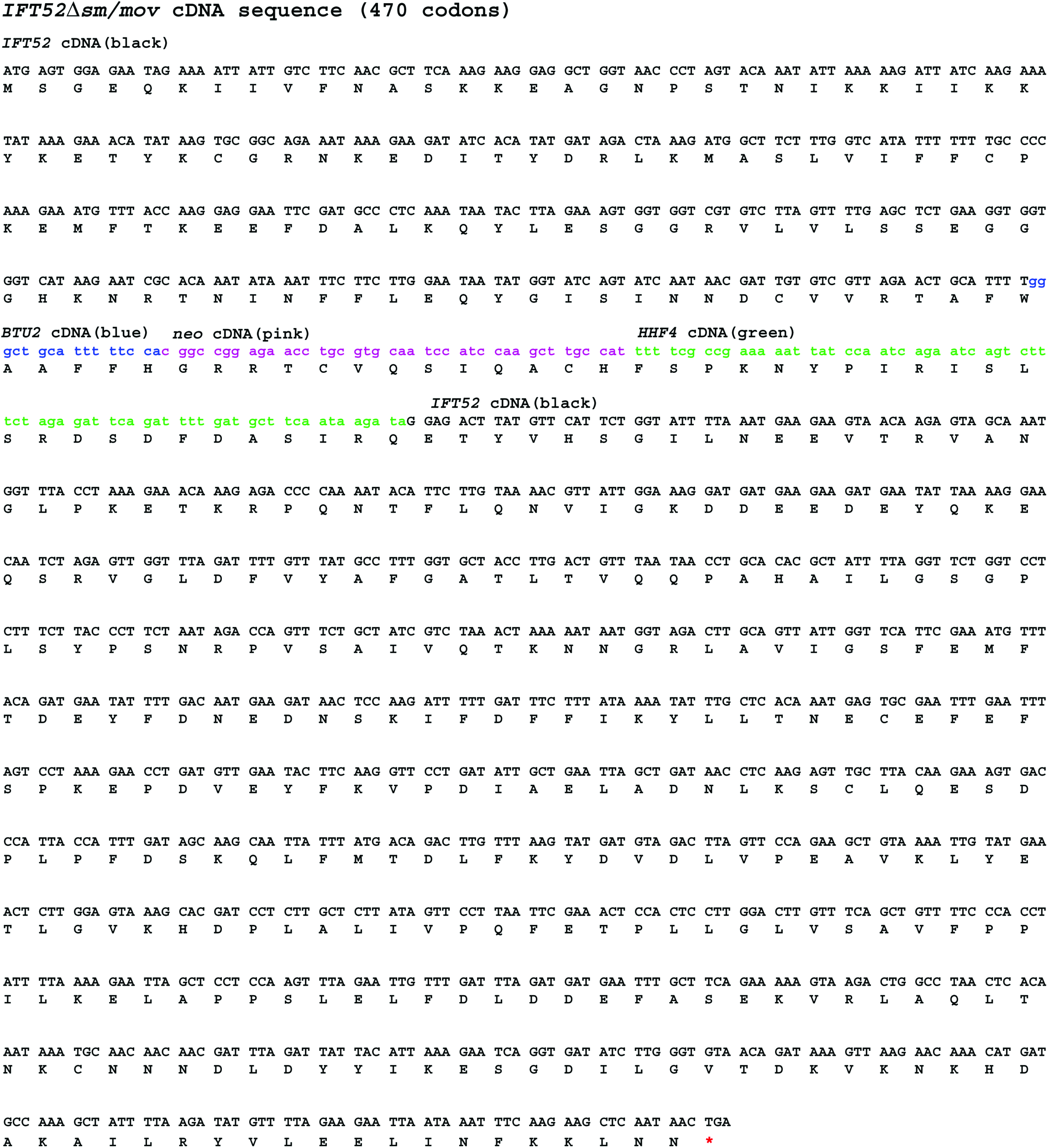
The cDNA sequence of *IFT52Δsm/mov* has 43 extra codons from residual *neo2* cassette. The residual *neo2* cassette consists of bacterial and *Tetrahymena* (*BTU2* and *HHF4*) sequences after being processed as artificial introns. The sequence shown corresponds to the region between exons 3 and 4 of the suppressed *IFT52* cDNA.

